# QuaC: A Pipeline Implementing Quality Control Best Practices for Genome Sequencing and Exome Sequencing Data

**DOI:** 10.1101/2023.03.06.531383

**Authors:** Manavalan Gajapathy, Brandon M. Wilk, Elizabeth A. Worthey

**Affiliations:** Center for Computational Genomics and Data Science, The University of Alabama at Birmingham, Birmingham, AL, USA; Department of Genetics, Heersink School of Medicine, The University of Alabama at Birmingham, Birmingham, AL, USA

## Abstract

Quality Control (QC) of human genome sequencing and exome sequencing data is necessary to ensure they are of sufficient quality for downstream analyses. While several QC tools are available to measure quality parameters at various levels post-sequencing, their output needs to be reviewed and interpreted in a very manual and time-consuming process. Such manual review is a major challenge towards standardization and consistency, as the process can be subjective depending on the reviewer. To address these difficulties, we have developed QuaC, which implements, integrates, and standardizes QC best practices at our Center. It performs three major steps: (1) runs several QC tools using data produced by the read alignment (BAM) and small variant calling (VCF) as input and optionally accepts QC output for raw sequencing reads (FASTQ); (2) executes QuaC-Watch to perform QC checkup based on the expected thresholds for quality metrics; and (3) aggregates QC metrics produced by all the QC tools as well as QuaC-Watch results into single, self-contained MultiQC report, both at the per-sample and across-project levels. This report provides aggregate summaries for all samples within a project/cohort for efficient comprehensive review while still allowing for granular review down to individual metrics for a single sample. Finally, we have developed a “Sample QC review system” schema to standardize QC reviewer’s logging of results and simplify downstream users’ interpretation of the reviewers finding.

## Statement of need

Application of Genome sequencing (GS) and exome sequencing (ES) based approaches has increased dramatically for both research and clinical purposes over the last decade. Several quality control (QC) tools have become available to help ensure that sequenced reads meet expected measures of quality, and to identify process related errors such as sample swaps or contamination. In recent years, efforts have been made to define QC metrics and acceptable thresholds for QC standardization across research groups (Kobren et al., 2021; Marshall et al., 2020). Despite these advances, integrating QC output from multiple tools, performing QC review in a standardized manner, and logging QC review results in an accessible and easy-to-understand manner to inform downstream consumers of the data remains a burden. Lack of defined procedures and appropriate shareable outputs for the latter step can result in downstream consumers proceeding unaware of QC issues. Without these outputs, downstream consumers often re-generate QC metrics, at times with limited expertise, wasting time and effort. Here, we present QuaC, a pipeline that integrates several QC tools and summarizes QC metrics for GS and ES samples using pre-defined and user-configurable thresholds to highlight potentially problematic samples. Further, we provide a system for interpretation of QC metrics called the “Sample QC Review System”, which supports recording of QC review results in a standardized manner.

## Quac Development

QuaC is a configurable pipeline developed using Snakemake and Python. QuaC provides a command-line interface (CLI), written in Python, to support user input, configuration, and execution. System-level tests along with mock data and example input configuration files are included in QuaC to assert correct operation after install and test future developments. Unit jobs triggered by QuaC are executed in Singularity container environment, as such setup provides the major advantage of reproducibility and portability across various user environments. QuaC is run at the project level, and samples in the project are provided as input in a pedigree file format (`.ped`), where sample metadata such as sample relatedness and sex can be optionally provided.

## QC Tools Utilized

QuaC runs several QC tools (Table 1) using BAM and VCF files as input. These support identification of sequencing, alignment, and variant calling related issues, within-species contamination, and sample swaps or incorrectly stated relationships between samples based on sex, ancestry, and relatedness estimations. Besides these tools, QuaC can optionally consume output from three QC tools executed separate from QuaC: FastQC to check quality of raw sequence reads (Andrews et al., 2012), FastQ Screen to check for cross-species contamination using raw sequence reads (Wingett & Andrews, 2018), and Picard-MarkDuplicates to check for read duplication in BAM files (*Picard Toolkit*, n.d.). While QuaC cannot run these QC tools, it can utilize their output as part of QC metric aggregation and summarization.

**Table 1:** QC tools used in QuaC. Note that this list does not include tools that QuaC can consume when run with `--include_prior_qc` flag.

| Tool | Usage in QuaC | QC type |
| --- | --- | --- |
| Qualimap (Okonechnikov et al., 2015) | Summarizes several alignment metrics using BAM file | BAM quality |
| Picard-CollectMultipleMetrics (Picard Toolkit, n.d.) | Summarizes alignment metrics from BAM file using several modules | BAM quality |
| Picard-CollectWgsMetrics (Picard Toolkit, n.d.) | Collects metrics about coverage and performance using BAM file | BAM quality |
| Mosdepth (Brent S. Pedersen & Quinlan, 2018) | Fast alignment depth calculation using BAM file | BAM quality |
| Indexcov (Brent S. Pedersen et al., 2017) | Estimate coverage from BAM index for GS (Skipped in exome mode) | BAM quality |
| Covviz (Covviz, n.d.) | Identifies large, coverage-based anomalies for GS using Indexcov output (Skipped in exome mode) | BAM quality |
| Bcftools stats (Danecek et al., 2021) | Summarizes VCF file stats | VCF quality |
| VerifyBamID2 (Zhang et al., 2020) | Estimates within-species (i.e., cross-sample) contamination using BAM file | Within-species contamination |
| Somalier (Brent S. Pedersen et al., 2020) | Estimation of sex, ancestry and relatedness using BAM file | Sex, ancestry, and relatedness estimation |

## QC Checkup Using QuaC-Watch

QuaC includes a tool called QuaC-Watch, which consumes results from the above-mentioned QC tools, compares QC metrics against the acceptable thresholds, and summarizes results using color-coded pass/fail flags for efficient review (Figure 1). This summary allows users to quickly review output from multiple QC tools, identify whether samples meet expected quality thresholds, and readily highlight samples that need further review. Reasonable default thresholds for QC metrics have been carefully selected and built in to QuaC-Watch. These are applicable for most GS and ES but are also configurable by the user. QC metrics and thresholds were curated based on literature (Kobren et al., 2021; Marshall et al., 2020), in-house analyses using many hundreds of both GS and ES samples, and knowledge gained from our past experiences. Integration of QC metrics and associated thresholds into QuaC not only assists with standardization of our internal QC review process, but also supports review and reusability between groups. We believe release of this information provides utility to the community. To our knowledge, this type of curated collection spanning an integrated suite of tools has not been made available previously.

**Figure 1:**
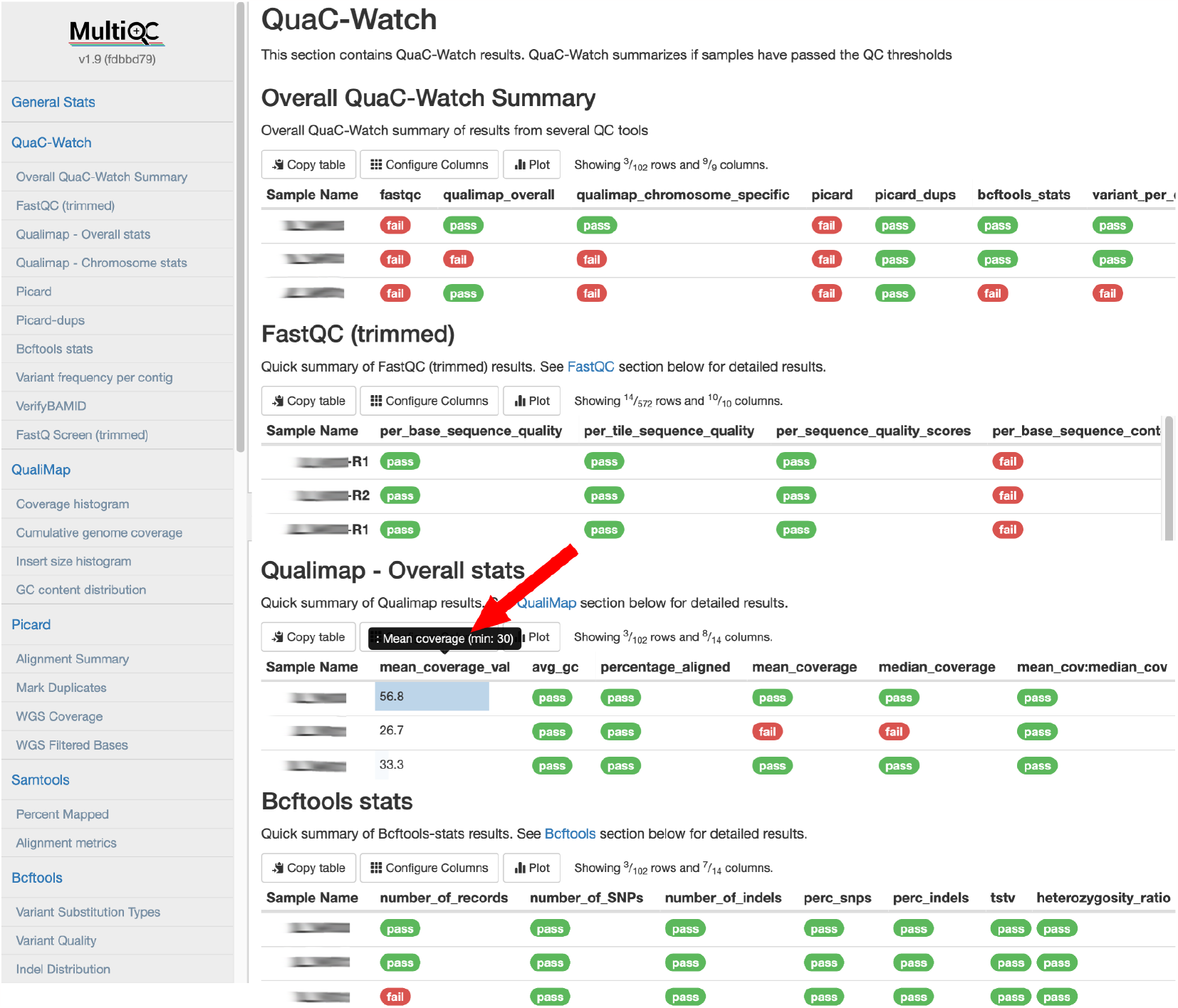
Aggregation and visualization of QC tools output and QuaC-Watch output using MultiQC at the project level. QuaC-Watch section shown here enables quick review of samples’ QC results and helps to quickly identify samples that need further review. Users may optionally toggle columns to view values for QC metrics of interest and hover over the column title to view thresholds used by QuaC-Watch (highlighted by red arrow). In addition to this project-level report, similar MultiQC report is created at the single-sample level for all the samples, which shows summarized QC results for only one sample..

## QC Aggregation

To minimize the time needed to review QC metrics and assess quality of samples across a project QuaC aggregates results produced by all the QC tools and QuaC-Watch, using MultiQC (Ewels et al., 2016), into per-sample and across-project stand-alone interactive HTML reports. The QuaC-Watch summary is presented as the first section of the report for initial review, followed by individual QC tool outputs for deeper review of metrics where high-level findings warrant it (Figure 1). Availability of MultiQC reports at both sample and project level enables easier review and distribution of QC results internally as well as with external collaborators.

## QC Review Process

Consistent and understandable dissemination of QC review results can be challenging when quality issues are identified, and even more so when these issues hamper accurate downstream analyses or interpretation. To reduce this burden, we devised a “Sample QC review system” where QC review results are flagged as pass, acceptable, poor, and fail, along with a free text field for review comments (Table 2). This system allows data consumers to rapidly review for sample issues and also points them to the known or likely cause of the issue. Since not all QC issues are catastrophic, this aids in rapid determination as to whether specific samples can be used for intended purposes. As not all users are proficient in interpreting results from the various QC tools, this system has proven helpful in enabling assessment and ensuring the quality of the conclusions based on this data.

**Table 2:** Fields logged in Sample QC database using controlled flags. Type 1 flags are pass, acceptable, poor, and fail. Type 2 flags are pass, fail, and not applicable.

| Field | Explanation | Allowed values |
| --- | --- | --- |
| Sample - Overall Status | Overall QC status considering results of all QC performed | Type 1 flags |
| FASTQ | Overall QC status considering results of all QC performed at FASTQ level | Type 1 flags |
| FASTQ Comment | Comments on QC at FASTQ level (e.g., small insert size, high adapter content, etc.) | Free text |
| BAM | Overall QC status considering results of all QC performed at BAM level | Type 1 flags |
| BAM Comment | Comments on QC at BAM level (e.g., low mean coverage, high duplication rate, etc.) | Free text |
| VCF | Overall QC status considering results of all QC performed at VCF level | Type 1 flags |
| VCF Comment | Comments on QC at VCF level (e.g., small insert size, high adapter content, etc.) | Free text |
| Other Species Contamination | Sample contamination status due to other species' genomic material | Type 1 flags |
| Human Cross-contamination | Sample contamination status due to other human's genomic material | Type 1 flags |
| Sex Check | Did the predicted sex match the expected sex? | Type 2 flags |
| Relatedness Check | Did the predicted relatedness match expected relatedness? | Type 2 flags |
| Ancestry Check | Did the predicted ancestry match expected ancestry? | Type 2 flags |
| Other Comments/Notes | Any other comments/notes concerning QC | Free text |

## Source Code and Documentation

Source code for QuaC is available for download at https://github.com/uab-cgds-worthey/quac under GNU GPLv3 license. Installation, setup, configuration, and usage documentation is available at https://quac.readthedocs.io.

## Acknowledgements

*We would like to thank Donna Brown for providing feedback on the utility of QuaC-Watch in research projects.*

*This work was supported in part by an award from the CF Foundation to Dr. Worthey (WORTHE19A0) and from UAB SOM Start-up funds to Dr. Worthey.*

